# The rostral medial frontal cortex is crucial for engagement in consummatory behavior

**DOI:** 10.1101/2022.03.11.484010

**Authors:** Samantha R. White, Mark Laubach

**Affiliations:** Department of Neuroscience, American University, Washington, DC, USA, 20016

**Author notes:** Corresponding author: Mark Laubach, PhD.

## Abstract

The medial frontal cortex (MFC) in rodents emits rhythmic activity that is entrained to the animal’s licking cycle during consumption and encodes the value of consumed fluids (Horst and Laubach, 2013; Amarante et al., 2017; Amarante and Laubach, 2021). These signals are especially prominent in the rostral half of the MFC. This region is located above an orbitofrontal region where mu opioid receptors regulate intake (Mena et al., 2011; Castro and Berridge, 2017) and reversible inactivation reduces behavioral measures associated with the incentive value and palatability of liquid sucrose (Parent et al., 2015a). Here, we examined the effects of reversible inactivation and stimulation of mu opioid receptors in rostral MFC on behavior in an incentive contrast licking task. Adult male rats licked to receive access to liquid sucrose, which alternated between high (16%) and low (4%) values over 30 sec periods. Bilateral infusion of muscimol reduced the total number of licks emitted over the 30 min test sessions, the time spent actively consuming sucrose, and the ratio of licks for the higher and lower value fluids. Inactivation did not alter licking frequency or variability or microstructural measures such as the duration of licking bouts that are classically associated with the palatability of a liquid reward. Infusions of DAMGO (1μg/μL) at the same sites had inconsistent behavioral effects across different subjects. Our findings suggest that the rostral MFC has a distinct role in the control of consummatory behavior and contributes to peristent consumption and not to the expression of palatability.

**SIGNIFICANCE STATEMENT:** The medial frontal cortex (MFC) of rodents has received attention in recent years and is considered as a singular cortical region with a potential unitary function. Increasing evidence suggests that MFC is composed of distinct subregions, with unique roles in the control of behavior. The present study adds to this literature by showing unique effects of reversibly inactivating the most rostral part of the medial frontal cortex and a lack of consistent effects of stimulating mu opioid receptors in the subregion. Findings are in contrast to previous reports on the more ventral orbitofrontal cortex and caudal medial frontal cortex and are important for understanding the general role of the rodent frontal cortex and how opioids may act to control behavior.

---

Information about rewarding stimuli is encoded in the rostral part of the medial frontal cortex (MFC), specifically in the rostral prelimbic area (aka area 32) and the laterally adjacent medial agranular cortex (Petykó et al., 2009; Horst and Laubach, 2013; Petykó et al., 2015; Amarante et al., 2017; Amarante and Laubach, 2021). Neurons in this region change firing rates around the initiation of sustained licking bouts (Petykó et al., 2009) and rhythmic neural activity, measured in field potentials and spike activity, is synchronized to the onset of consummatory behavior (Horst and Laubach, 2013). Using an incentive contrast licking task using liquid sucrose rewards (Flaherty, 1999; Dwyer, 2012), Amarante et al. (2017) found that consummatory-related activity in rostral MFC develops with experience, reflects differences in reward value, and is reduced by inactivation of the rostral MFC in the opposite hemisphere. In a related study, Amarante and Laubach (2021) found that rhythmic activity during consumption is directionally coherent with the timing of rhythms in the rostral MFC leading those in the lateral orbitofrontal cortex (OFC) and further reflects differences in value for sucrose concentration and fluid volume.

The rostral MFC region examined in the studies described above has not been studied using bilateral reversible inactivation with muscimol, a GABA-A agonist (Figure 1). Therefore, the role of the region in the control of consummatory behavior is not known. Inactivations in more ventral parts of the frontal cortex using the incentive contrast licking task were found to disrupt the expression of incentive contrast and behaviors associated with palatability (Parent et al., 2015a). It is not known if dorsal parts of the rostral MFC have a role in incentive contrast, palatability, or active engagement in consumption (Swanson et al., 2019). One of the goals of the present study was to determine how reversible inactivations of the rostral MFC alter consummatory behavior.

**Figure 1:**
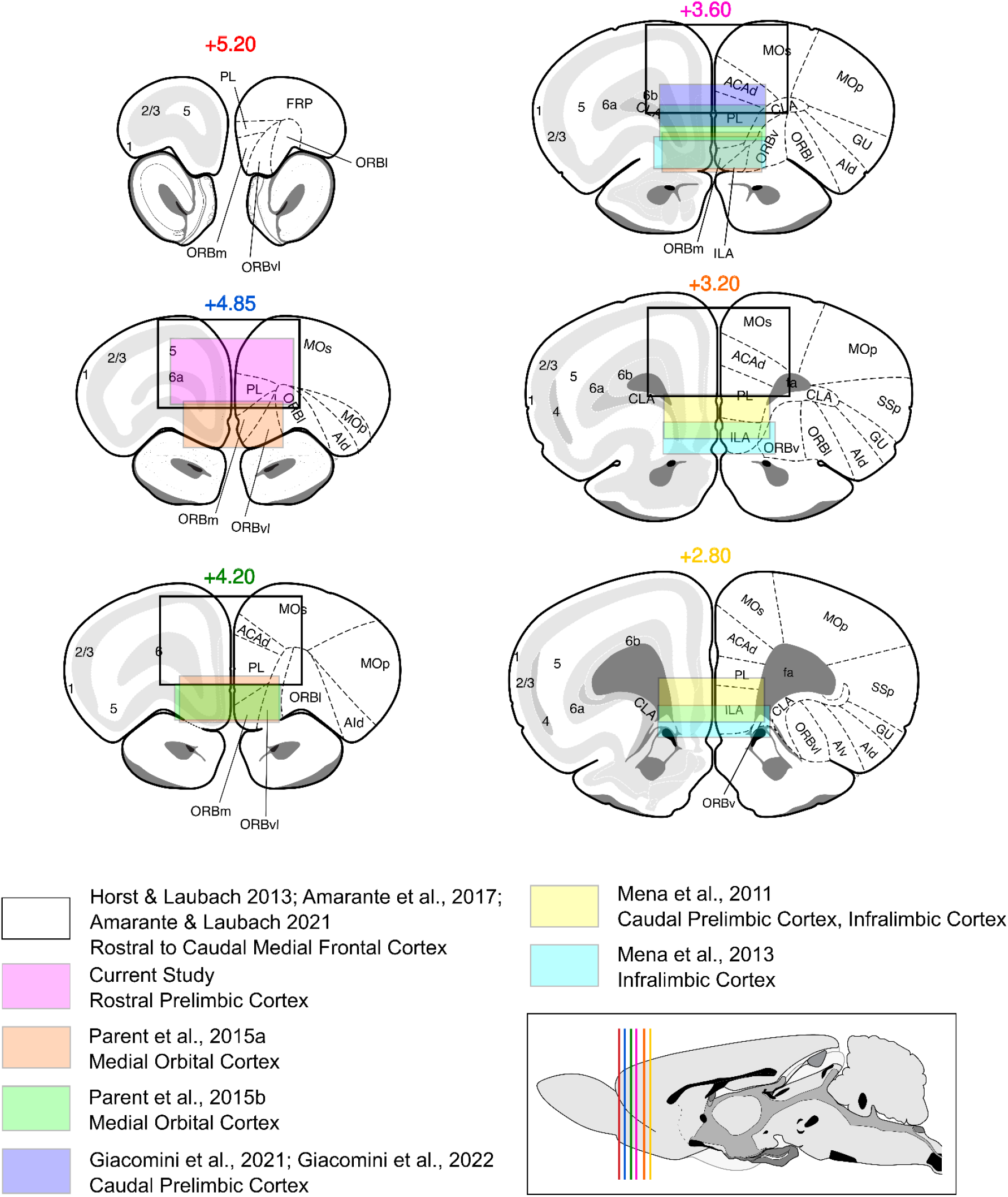
Frontal cortical anatomy for the context of this study. Previous work in the medial frontal cortex spans a rostral to caudal axis, as well as a dorsal and ventral axis. This figure depicts the locations of previous studies on this cortical region and adjacent regions that used consummatory tasks. Neural recordings by Horst & Laubach (2013), Amarante et al. (2017), and Amarante & Laubach (2021) (black outlines) were made in the rostral medial frontal cortex, in the prelimbic and medial agranular cortices. Parent et al. (2015a,b; orange, green) targeted ventral portions of rostral MFC, considered to be medial orbital cortex. Mena et al. (2011) (yellow) targeted caudal MFC and were more ventral on the medial wall, at the border between the prelimbic and infralimbic cortices. Mena et al. (2013) (light blue) targeted the infralimbic cortex. Giacomini et al. (2021, 2022; dark blue) studied differences across multiple regions of the frontal cortex, but prelimbic sites were restricted to the caudal part of the prelimbic cortex. The current study (pink) targeted the rostral prelimbic cortex. Colored boxes highlight the regions associated with different studies. The left side of each coronal slice represents the laminar organization of the cortex according to the Swanson Atlas. The right side of each slice represents area delineations of cortical regions according to the Swanson Atlas and Paxinos Atlas. Abbreviations, according to the Swanson Atlas: ACAd: anterior cingulate area, dorsal part, AId: agranular insular area, dorsal part, AIv: agranular insular area, ventral part, CLA: claustrum, fa: corpus callosum, anterior forceps, FRP: frontal pole, cerebral cortex, GU: gustatory area, ILA: infralimbic area, MOp: primary somatomotor area, MOs: secondary somatomotor areas, ORBl: orbital area, lateral part, ORBm: orbital area, medial part, ORBv: orbital area, ventral part, ORBvl: orbital area, ventrolateral part, PIR: piriform area, PL: prelimbic area, SSp: primary somatosensory area, SSp-m: primary somatosensory area, mouth region.

There has been a recent interest in understanding the roles of various neurotransmitter systems in modulating reward processing in the frontal cortex. Opioid receptors in the frontal cortex have been of special interst in recent years, given the ongoing opioid crisis. Studies on the effects of the selective mu opioid agonist DAMGO across various regions of the frontal cortex have reported effects on food intake, patterns of feeding, and overall activity (Mena et al., 2011, 2013; Selleck et al., 2018; Giacomini et al., 2021, 2022). Castro and Berridge (2017) reported evidence for opioid and orexin “hot spots” in the frontal cortex, including in the rostral MFC, where infusions of DAMGO or orexin induce changes in the expression of the immediate early gene Fos and have corresponding effects on hedonic orofacial reactions and changes in feeding. None of these studies reported pharmacologyical data from the dorsal part of the most rostral MFC. Therefore, in the present study we also examined how local stimulation of mu opioid receptors might alter behavior in the incentive contrast licking task.

Anatomical studies have established that neurons in the rostral MFC project to brainstem nuclei involved in jaw movements (Yoshida et al., 2009). These connections do not exist for the more ventral prelimbic and medial orbital regions studied by Parent et al. (2015a) (e.g. Gabbott et al., 2003; Hoover and Vertes, 2007). If rostral MFC neurons are involved in modulating brainstem nuclei controlling jaw movements, then perturbations of rostral MFC might have sensorimotor effects on consummatory behavior, such as interlick intervals (e.g. Gaffield and Christie, 2017) or the ability to engage in licking. We found that reversible inactivation of the rostral MFC has distinct effects compared to inactivating the ventrally located medial OFC. Rather than affecting measures of palatability, inactivation reduced engagement in consumption without changing sensorimotor measures. The effects of rostral MFC inactivation might best be described in terms of abulia (Berrios and Gili, 1995), a reduction in the expression of “willful” and goal-directed movements. Infusions of DAMGO had no consistent effect on any behavioral measures across rats.

## MATERIALS AND METHODS

All procedures were approved by the Animal Care and Use Committee at American University (Washington, DC). Procedures conformed to the standards of the National Institutes of Health Guide for the Care and Use of Laboratory Animals. All efforts were taken to minimize the number of animals used and to reduce pain and suffering.

### Animals

Ten male rats (seven Sprague-Dawley (Charles River), three Long Evans (Envigo)) of 400-450 grams were used in this study. Animals were housed individually and kept on a 12/12 h light/dark cycle switching at 7:00 AM and 7:00 PM. Upon arrival, animals were given one week of habituation to their new environment with free access to rat chow followed by daily handling for one week. After habituation and initial daily handling, animals had regulated access to food to maintain their body weights at approximately 90% of their free-access weights. Rats typically received 14–18 g of food each day around 5 pm and were weighed daily throughout the period of training and testing in an incentive contrast licking task. Animals had free access to water throughout the experiments. Of the rats used in this study, one rat was removed due to an unintentional unilateral infusion from a blockage in the guide cannula.

### Behavioral Apparatus

All animals were trained in sound-attenuating behavioral boxes (ENV-008; Med Associates) containing a single horizontally placed spout located on one wall at 6.5 cm from the floor and a house light at the top of the box. Control of pumps and behavioral quantification was done using a MedPC system version IV (Med Associates). The licking spout was custom built to allow the convergence of two independent solution lines stemming from two independent pumps at a single point. Licking was tracked optically as breakage of an infrared beam by the tongue between a custom built emitter/detector placed directly in front of the licking spout. Solution lines were connected to 60cc syringes and solution was made available to animals by lick-triggered, single speed pumps (PHM-100; Med Associates) which drove syringe plungers. Each lick activated a pump which delivered roughly 30 μL per 0.5 second activation.

### Behavioral Task

After rats could produce >700 licks for 16% sucrose in a single session, they were conditioned in the incentive contrast licking task (Figure 2A). The task used in these experiments is the same as described previously (Parent et al., 2015a; Amarante et al., 2017). Briefly, animals were placed into the operant chamber for 30 min and had constant access to the spout. Two independent pumps delivered sucrose solution to the same spout and were loaded with syringes containing either high value 16% (wt/vol) sucrose solution or low value 4% (wt/vol) sucrose solution. Licking at the spout initiated a 30-s epoch of access to the high value solution. Each lick was recorded and a lick occurring after the end of the 30-s epoch triggered a 30-s epoch of access of low value sucrose. These epochs of access continually switched back and forth between pumps and provided alternating access to high and low value solutions. At the end of the 30 min session, the house light turned off and animals stopped receiving sucrose solution. Quantification of behavior was implemented via analysis of metrics for consumption (time on task, total licks, and proportion of high value licks) and metrics of licking microstructure (duration of licking bouts, number of licking bouts, and intra-bout licking rates).

**Figure 2:**
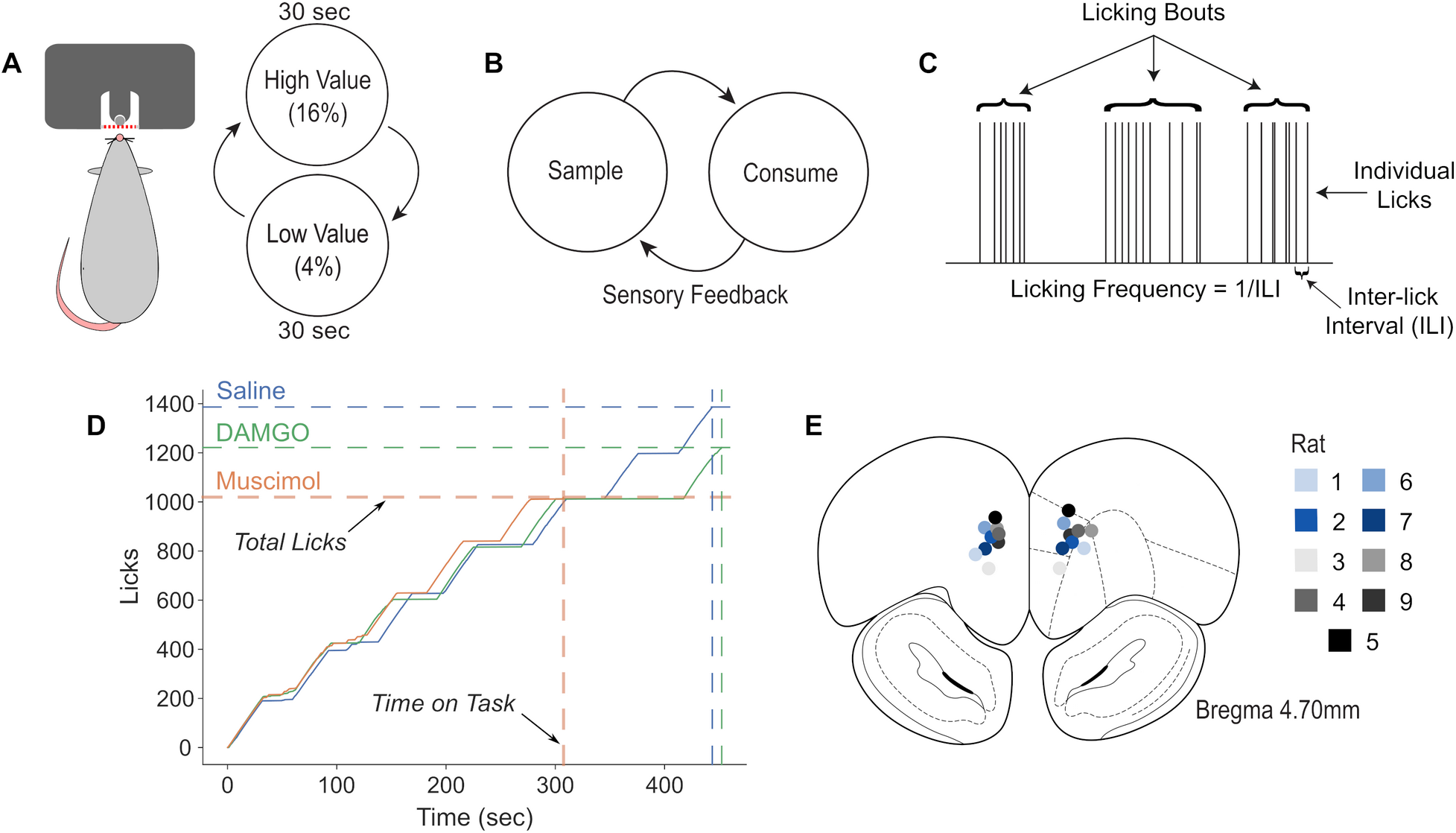
Task, Lick Microstructure, and Cannula Placement. **(A)** Rats licked on a single spout and received access to solutions containing relatively high (16% wt/vol) or low (4%) sucrose. The two fluids were presented in alternating 30-s epochs at the same spout. (**B**) Conceptual model of task. Rats lick to sample fluid during initial learning and later when low value reward is available. They show increased licking for the higher value reward over the first several days of training (Figure 1C in Parent et al., 2015a). They transition between sampling and consuming after initial training. Shifts between the states is guided by sensory feedback (taste/flavor of fluid). The proportion of high value licks reflects consummatory licking. **(C)** Animals consumed sucrose by emitting bouts of licks. Key measures of licking microstructure are indicated. **(D)** Total Licks and Time on Task are depicted from a session using a cumulative record plot. The data are from the rat with the median value for Time On Task. **(E)** Histological confirmation revealed that the cannula across the nine subjects were consistent in the rostral medial frontal cortex (+4.7 AP, +/-0.7 ML, and −2.0 DV).

### Surgery

Prior to cannulation, animals were given 2-3 days of free access to rat chow and water. Anesthesia was induced by an injection of diazepam (5 mg/kg, IP) and maintained with isoflurane (3.0%; flow rate 5.0 cc/min). The scalp was shaved clean and animals were injected with a bolus of carprofen subcutaneously. Animals were placed into a stereotaxic frame using non-penetrating ear bars and the skull was covered with iodine for 1 min. Iodine was wiped clean from the scalp and the eyes were covered with ophthalmic ointment to prevent drying over the span of the surgery. Lidocaine (0.5 mL) was injected under the scalp and an incision was made longitudinally along the skull. The skin was retracted laterally and all tissue was cleaned from the surface of the scalp. The skull was leveled by adjusting the stereotaxic apparatus to ensure bregma and lambda were within the same horizontal plane. Four skull screws were placed along the edges of the skull for support and adhesion of the implant to the skull. Craniotomies were drilled bilaterally in the frontal skull plates over the rostral medial frontal cortex and 26 gauge stainless steel guide cannula (Plastics One) were lowered into rostral MFC (coordinates from bregma AP +3.2mm, ML +/1.2mm, DV −2.0mm surface of the brain at a 12-degree posterior angle) (Paxinos and Watson, 2014). The guide cannula contained a 33 gauge stainless steel wire (“dummy cannula”) that extended 0.4mm past the tip of the guide cannula. Craniotomies were sealed using cyanoacrylate (Slo-Zap) and an accelerator (Zip Kicker), and methyl methacrylate dental cement (AM Systems) was applied and affixed to the skull via the skull screws. Animals were given Carprofen (5 mg/kg, s.c.) for postoperative analgesia. Animals recovered from surgery in their home cages for at least 1 week with full food and water, and were weighed and monitored daily for 1 week after surgery. After 1 week, animals’ body weights returned to presurgical levels, restricted access to rat chow was reinstated, and animals continued with daily behavioral testing sessions.

### Drug Infusions

Following recovery from surgery (5-7 days) and a period of one week reacclimation to the task with restricted food access, a series of controls were performed on all rats. First, animals were exposed to the same duration and levels of isoflurane gas used during drug infusions to control for exposure to isoflurane. Second, a PBS control was carried out where the same volume of vehicle without drug was infused into the rostral MFC while the animals were anesthetized. Finally, on test day, animals were anesthetized via isoflurane and drug was infused centrally into the rostral MFC. Following test day, recovery sessions were carried out. Each rat received between 3-4 sessions of drug infusions during the time of this study. Drugs used in this study included muscimol (1.0μg/μL) and DAMGO ([d-Ala2, N-Me-Phe4, Gly5-ol]-enkephalin) (1.0μg/μL). All drugs were obtained from Tocris and made into solutions using sterile PBS with pH 7.4. Concentrations were based on published studies (Parent et al., 2015a; Giacomini et al., 2021).

Infusions were performed by inserting a 33-gauge injector into the guide cannula. The injector extended 0.4mm past the tip of the guide cannula. A volume of 1.0uL of fluid (Parent et al., 2015a) was delivered at a rate of 0.25uL/min with a syringe infusion pump (KDS Scientific). The injector was connected to a 10 uL Hamilton syringe via 0.38 mm diameter polyethylene tubing. After infusion was finished, the injector was left in place for at least 2 minutes to allow for diffusion of the fluid. The injector was slowly removed and the dummy cannula was replaced. Rats were tested in the incentive contrast licking task 1 hour after the PBS and muscimol infusions or 20-30 minutes after DAMGO infusions.

### Statistical analysis

All data were analyzed using Python (Anaconda distribution: https://www.continuum.io/). Analyses were run as Jupyter notebooks (http://jupyter.org/). Statistical testing was performed using DABEST, a Python package for data analysis using bootstrap-coupled estimation. Two-sided, paired permutation tests were used to compare saline and drug infusion conditions (Ho et al., 2019). 5000 bootstrap samples were taken; the confidence interval is bias-corrected and accelerated. The p-values reported are the likelihood of observing the effect size if the null hypothesis of zero difference is true. For each p-value, 5000 reshuffles of the control and test labels were performed. Results are displayed as estimation plots produced by DABEST. They present the raw data and the bootstrap confidence interval of the effect size (the difference in means) as a single integrated plot.

### Behavioral Data Analysis

Primary measures are described in Figure 2C. Analysis of licking used custom scripts written in Python. Inter-lick intervals greater than 1 s or less than 0.09 s were excluded from the analysis. Detection and quantification of licking bouts were done as in previous studies (Gutierrez et al., 2010; Horst and Laubach, 2013; Parent et al., 2015a). Specifically, bouts were defined as having at least three licks within 300 ms and with an interbout interval of 1s or longer. Behavioral measures included time on task (the total time rats spent engaged in sustained licking), total licks across the session, proportion of high value licks out of the total licks, the duration and number of licking bouts, and the lick frequency (inverse of the median inter-lick interval).

### Confirmation of Cannula Placement

At the termination of experiments, animals were initially anesthetized with isoflurane gas and injected intraperitoneally with Euthasol. Animals were transcardially perfused first with 500 ml of cold saline solution followed by 500 ml of cold 4% paraformaldehyde. Brains were removed and post-fixed in a mixture containing 4% paraformaldehyde, 20% sucrose, and 20% glycerol. Brains were then cut into 100 μm-thick coronal slices using a freezing microtome. Brain sections were mounted onto gelatin-coated slides and Nissl stained via treatment with thionin. Thionin-treated slices were dried through a series of alcohol steps and cleared with Xylene. Slides were covered with permount and coverslipped. Sections were imaged using a Tritech Research scope (BX-51-F), Moticam Pro 282B camera, and Motic Images Plus 2.0 software. The most ventral point of the injection bolus was compared against the Paxinos and Watson atlas to confirm coordinates.

## RESULTS

### Anatomical definition of the rostral medial frontal cortex

This study focused on a rostral region of the MFC that has not been previously examined using bilateral reversible inactivation or intra-cortical pharmacology (Figure 1). The prelimbic cortex spans from AP +5.20 to +2.50 mm relative to Bregma. At the most rostral limit, ahead of AP +5.20, the dorsal medial cortex is considered as the frontal pole and is characterized by a lack of a clear layer 6 (Swanson, 2004). Regions of MFC caudal to +4.80 but ahead of 3.60 is defined here as rostral MFC based on the following criteria: (i) no identifiable anterior forceps of the corpus callosum, (ii) six identifiable cortical layers, and (iii) the rhinal fissure is complete or nearly complete. The caudal MFC is posterior to the rostral MFC. Anatomical sections at this level of the frontal cortex show the following criteria: (i) presence of the anterior forceps of the corpus callosum, (ii) a clearly developed layer 6, and (iii) and a partial rhinal fissure.

### Inactivation of rostral MFC reduces engagement in consumption

Three behavioral measures associated with consummatory engagement were affected by bilateral reversible inactivation of the rostral MFC (Figure 2C). Muscimol reduced the overall time rats spent licking in the task (Figure 3A; paired mean difference: - 49.1; p-value: 0.022; 95%CI: −82.2, −19.6). Inactivation also reduced the total number of licks rats emitted across the testing sessions (Figure 3B; paired mean difference: −335.0; p-value: 0.0196; 95%CI: −576.0, −138.0). Finally, inactivation reduced the proportion of high value licks (Figure 3C; paired mean difference: −0.0425; p-value: 0.0396; 95%CI: −0.0619, 0.0114). Table 1 demonstrates that the data points where there was an opposite effect of muscimol for individual rats (increases rather than decreases in beahvioral measures and vice versa) are not driven by a singular rat.

**Figure 3:**
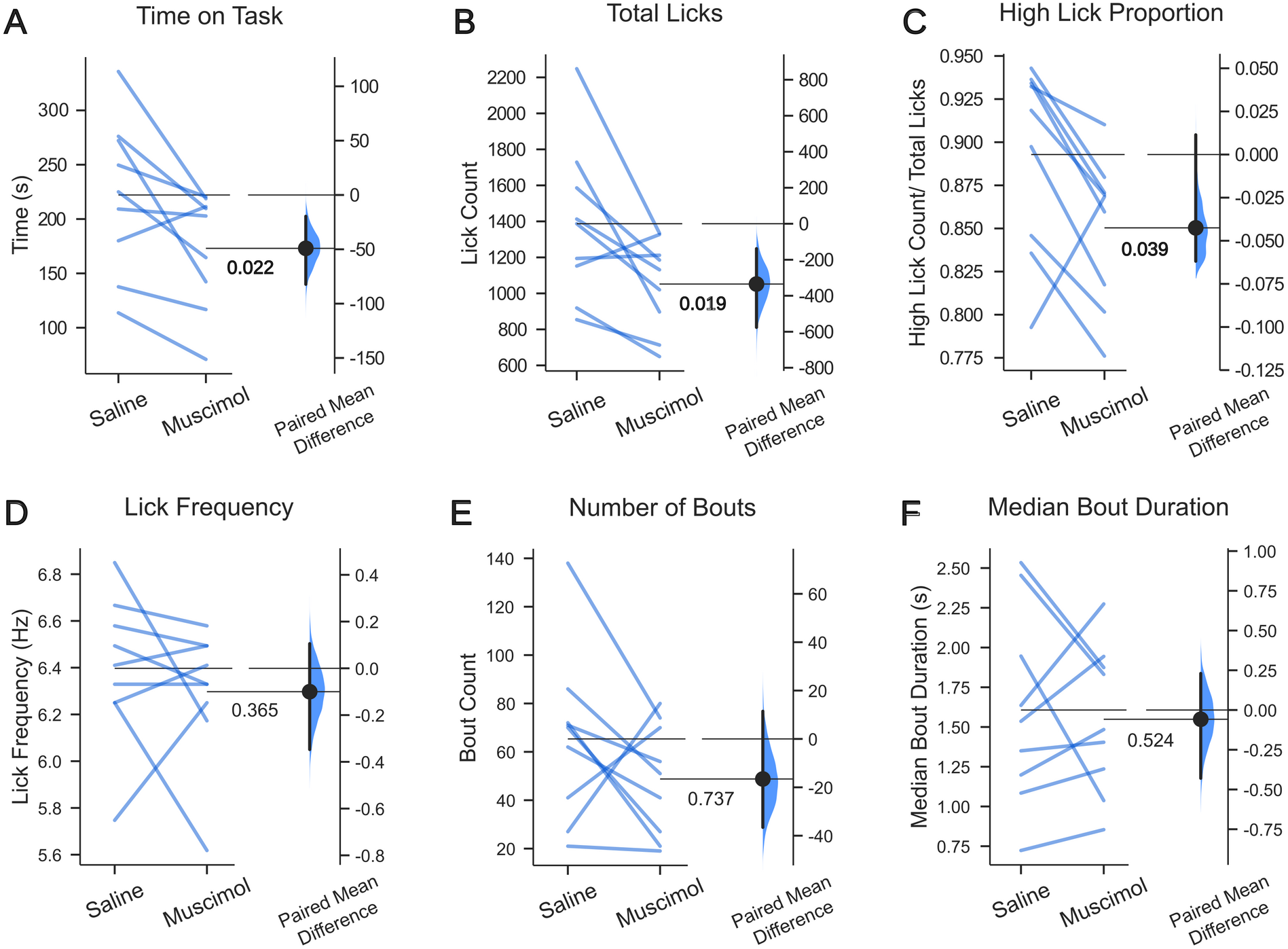
Inactivation of rostral MFC reduces task engagement in an incentive contrast licking task. However, inactivation had no effects on sensorimotor function or fluid palatability. Bilateral infusion of muscimol reduced **(A)** the time spent engaged in the task, **(B)** the total number of licks emitted over the 30 min test sessions, and **(C)** the proportion of licks for the high value fluid. Inactivation did not alter **(D)** licking frequency or microstructural licking measures such as **(E)** number of licking bouts emitted or **(F)** the duration of licking bouts, measures that are classically associated with the palatability of a liquid reward. Estimation plots of the values comparing saline to muscimol. The raw values are plotted as Tufte slopegraphs on the top portion of the figure; each line represents a single rat’s data between two conditions. The bottom portion plots the paired mean difference and a visual representation of the bootstrap confidence interval of the effect size. P-values displayed on each subpanel reflect the result of a two-sided permutation test (5000 resamples).

**Table 1:** Behavioral measures for each rat.

| Saline |  |  |  |  | Bouts |  |
| --- | --- | --- | --- | --- | --- | --- |
| Rat | Time On Task | Licks | High Prop. | Lick Freq. | Duration | Number |
| 1 | 209.228 | 1194 | 0.943 | 5.747 | 1.350 | 70 |
| 2 | 272.230 | 1729 | 0.846 | 6.410 | 1.536 | 72 |
| 3 | 335.774 | 2248 | 0.932 | 6.667 | 1.198 | 62 |
| 4 | 249.616 | 1413 | 0.934 | 6.250 | 2.454 | 41 |
| 5 | 180.006 | 1152 | 0.897 | 6.849 | 1.946 | 27 |
| 6 | 276.124 | 1586 | 0.918 | 6.329 | 2.534 | 71 |
| 7 | 225.014 | 1387 | 0.936 | 6.494 | 1.636 | 21 |
| 8 | 113.674 | 919 | 0.836 | 6.579 | 0.723 | 138 |
| 9 | 137.660 | 854 | 0.793 | 6.250 | 1.084 | 86 |

| Muscimol (1 µg/µl) |  |  |  |  | Bouts |  |
| --- | --- | --- | --- | --- | --- | --- |
| Rat | Time On Task | Licks | High Prop. | Lick Freq. | Duration | Number |
| 1 | 202.800 | 1212 | 0.880 | 6.250 | 1.404 | 27 |
| 2 | 142.262 | 897 | 0.802 | 6.494 | 1.944 | 21 |
| 3 | 218.706 | 1334 | 0.910 | 6.579 | 1.484 | 41 |
| 4 | 220.484 | 1131 | 0.860 | 5.618 | 1.831 | 70 |
| 5 | 211.394 | 1327 | 0.817 | 6.173 | 1.037 | 80 |
| 6 | 209.626 | 1185 | 0.869 | 6.329 | 1.874 | 56 |
| 7 | 164.530 | 1020 | 0.871 | 6.329 | 2.274 | 19 |
| 8 | 70.880 | 649 | 0.776 | 6.494 | 0.854 | 74 |
| 9 | 116.720 | 713 | 0.868 | 6.410 | 1.236 | 51 |

**Table 1: Behavioral measures for each rat.**
| DAMGO (1 µg/µl) |  |  |  |  | Bouts |  |
| --- | --- | --- | --- | --- | --- | --- |
| Rat | Time On Task | Licks | High Prop. | Lick Freq. | Duration | Number |
| 1 | 243.638 | 1429 | 0.919 | 6.250 | 1.186 | 80 |
| 2 | 197.926 | 1260 | 0.849 | 6.494 | 1.422 | 67 |
| 3 | 197.850 | 1289 | 0.877 | 6.757 | 1.030 | 71 |
| 4 | 340.336 | 1986 | 0.937 | 6.250 | 1.640 | 53 |
| 5 | 337.970 | 1948 | 0.755 | 6.329 | 1.606 | 81 |
| 6 | 185.088 | 1105 | 0.882 | 6.329 | 0.965 | 58 |
| 7 | 192.458 | 1222 | 0.909 | 6.579 | 3.708 | 22 |
| 8 | 124.444 | 944 | 0.863 | 6.494 | 0.732 | 151 |
| 9 | 107.620 | 655 | 0.791 | 6.098 | 1.052 | 57 |

### Inactivation of rostral MFC did not alter palatability or sensorimotor behavior

Bilateral infusion of muscimol in the rostral MFC had no effect on the rate at which rats licked for sucrose, the inverse of the median inter-lick interval (Figure 3D; paired mean difference: −0.1; p-value: 0.365; 95%CI: −0.347, 0.105). Furthermore, inactivation of rostral MFC had no effect on the number of bouts (lick clusters) across the session (Figure 3E; paired mean difference: −0.0581; p-value: 0.737; 95%CI: −0.429, 0.23) or the duration of licking bouts (Figure 3F; paired mean difference: −16.6; p-value: 0.225; 95%CI: −36.6, 11.4).

### Stimulation of mu opioid receptors had no consistent effects on behavior

Animals received infusions of the mu opioid receptor agonist DAMGO at the same sites were muscimol was infused. In contrast to the results of inactivation reported above, infusions of DAMGO had no consistent effects on any behavioral measure in the incentive contrast licking task. There were no group-level effects observed for the total time rats spent licking (Figure 4A; paired mean difference: −8.0; p-value: 0.807; 95%CI: −59.2, 53.9), the total number of licks emitted across the testing sessions (Figure 4B; paired mean difference: −71.6; p-value: 0.694; 95%CI: −395.0, 264.0), and the relative proportions of higher and lower value licks (Figure 4C; paired mean difference: −0.0282; p-value: 0.108; 95%CI: −0.0747, −0.00624). Infusions of DAMGO also had no consistent effects on inter-lick intervals (Figure 4D; paired mean difference: 0.000397; p-value: 1.0; 95%CI: −0.162, 0.16), the number of bouts (Figure 4E; paired mean difference: −0.124; p-value: 0.68; 95%CI: - 0.586, 0.62), and the durations of licking bouts (Figure 4F; paired mean difference: 5.78; p-value: 0.524; 95%CI: −6.44, 22.8). Table 1 demonstrates that the data points where there were variable effects for individual rats (increases rather than decreases in beahvioral measures and vice versa) are not driven by a singular rat or by a pattern of baseline behavior. There was also no evidence that variability in behavioral pattern could be explained by differences in precise anatomical location with respect to any axis (dorsal/ventral, medial/lateral, anterior/posterior) in the targeted rostral prelimbic cortex.

**Figure 4:**
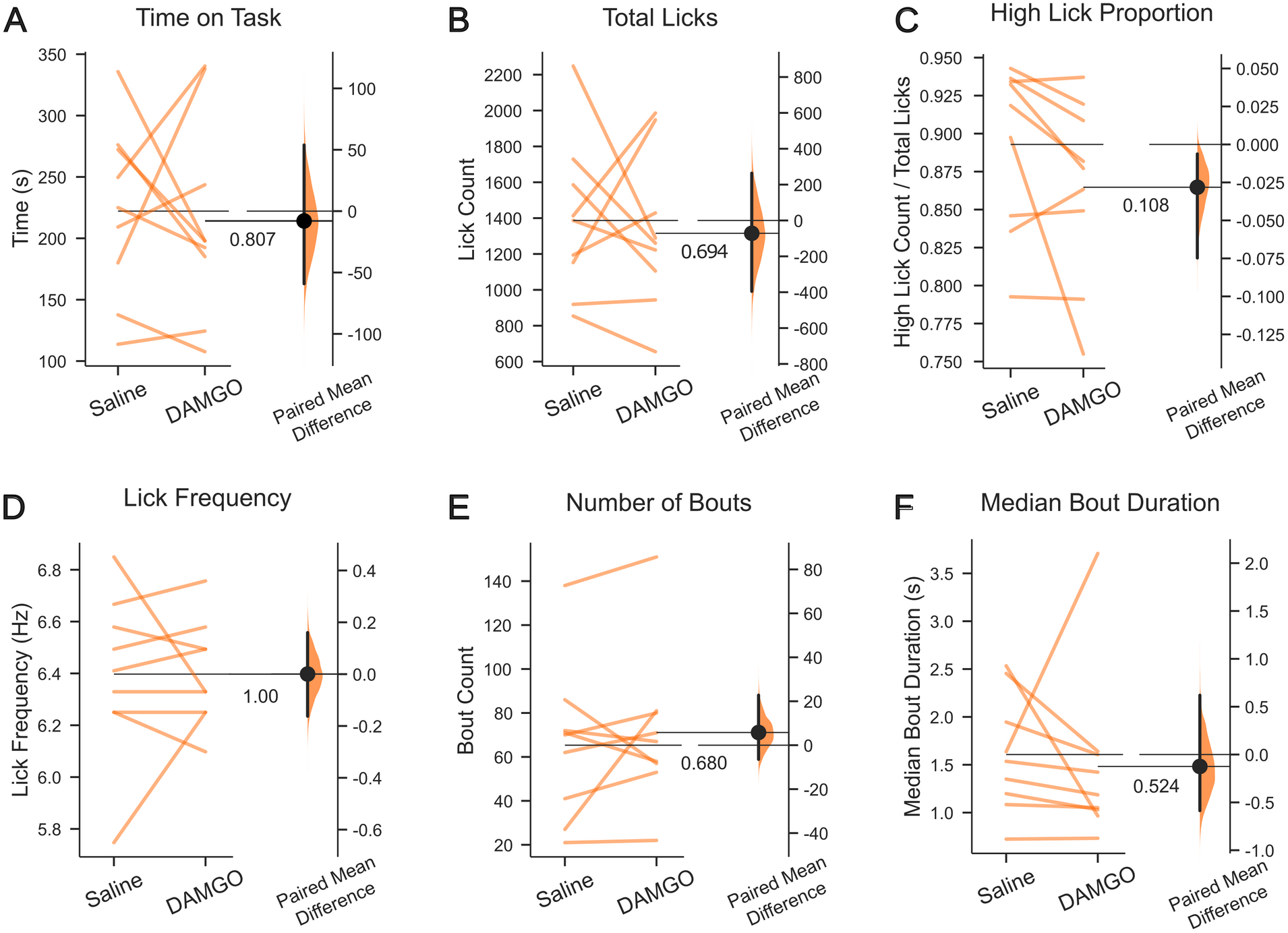
Activation of mu-opioid receptor in rostral MFC has no effect on task engagement, sensorimotor function or fluid palatability in an incentive contrast licking task. Bilateral infusion of DAMGO didn’t impact **(A)** the time spent engaged in the task, **(B)** the total number of licks emitted over the 30 min test sessions, or **(C)** the proportion of licks for the high value fluid. Further, it did not alter **(D)** licking frequency or microstructural licking measures such as **(E)** number of licking bouts emitted or **(F)** the duration of licking bouts, measures that are classically associated with the palatability of a liquid reward. Estimation plots of the values comparing saline to muscimol. The raw values are plotted as Tufte slopegraphs on the top portion of the figure; each line represents a single rat’s data between two conditions. The bottom portion plots the paired mean difference and a visual representation of the bootstrap confidence interval of the effect size. P-values displayed on each subpanel reflect the result of a two-sided permutation test (5000 resamples).

## DISCUSSION

In the current study, we sought to examine the role of the rostral medial frontal cortex (Figure 1) in an incentive contrast licking task via reversible inactivation and stimulation of mu opioid receptors. Adult male rats licked to receive access to liquid sucrose alternating between high (16%) and low (4%) values over 30 second periods, and learned to lick persistently when the higher value fluid was available. Bilateral infusion of muscimol reduced consummatory engagement, including the time spent licking, the total number of licks per session, and the proportion of licks for the high value fluid (Figure 3A-C). By contrast, infusions of DAMGO (1μg/μL) at the same locations had no consistent effects on any measures of task performance across rats (Figure 4A-C). Importantly, inactivation via muscimol and stimulation of mu receptors by DAMGO did not alter measures of gross sensorimotor function or fluid palatability, including licking frequency and the number and duration of sustained licking bouts (Figure 3D-F, Figure 4D-F).

The data reported here are important for understanding several electrophysiology studies on the rostral MFC (Horst and Laubach, 2013; Amarante et al., 2017; Amarante and Laubach, 2021). Horst and Laubach (2013) reported the highest proportion of recording sites that show phase locking around bouts of licking are within the rostral prelimbic sites (ahead of AP +3.6). Amarante et al. (2017) reported that neural activity in the rostral medial frontal cortex further encodes the value of liquid sucrose and a related study by Amarante and Laubach (2021) further showed that rostral MFC activity is synchronized with, and may even drive variability in, rhythmic activity in the lateral orbitofrontal cortex (Amarante and Laubach, 2021). These studies assumed that inactivation of the rostral MFC would disrupt licking microstructure and measures of palatability, based on studies described below that focused on the more ventral medial orbital cortex (Parent et al., 2015a). Reversible inactivations carried out in Amarante et al. (2017) were done unilaterally, to examine effects of rostral MFC activity on neural signals in the opposite hemisphere. The unilateral infusions in that study had no major effects on behavior, which was important to rule out effects in the opposite hemisphere due to changes in behavior. The present results establish that rather than mediating effects on palatability and incentive contrast, the neural signals reported by Amarante et al. (2017) and Amarante and Laubach (2021) reflected the animals’ engagement in the licking task, a process that could reflect abulia (a lack of willful or decisive action) or apathy, and very much supports the interpretation in the Amarante et al. (2017) and Amarante and Laubach (2021) studies of neural activity in the rostral MFC as being sensitive to “response vigor”.

The findings of this study are also important for understanding the roles of various parts of the rostral MFC in the control of consummatory behaviors. Pharmacological perturbations of the dorsal rostral MFC was found to be distinct from findings in nearby cortical areas such as the medial orbital (Parent et al., 2015a, 2015b), and ventral orbital,caudal prelimbic and infralimbic cortices (Giacomini et al 2022). In the studies by Parent and colleagues, inactivation of the medial orbital cortex (which was referred to by the authors as rostral medial prefrontal cortex) resulted in dramatically fragmented licking behavior, affecting bout duration and the variability of inter-lick intervals, and also disrupted the expression of incentive contrast. As shown in Figure 3 of Parent et al. (2015a) and Figure 2 of Parent et al. (2015b), most infusions of muscimol and other drugs were ventral to the locations in the present study. The medial orbital region impacts measures of palatability and sensorimotor behavior. By contrast, inactivation of the rostral prelimbic area, studied here, altered how long the rats were engaged in consummatory behavior (time spent licking and total licks per session) and not measures associated with palatability or sensorimotor function. These findings are also distinct from inactivation of caudal regions of MFC in a study by Giacomini et al (2022). They reported that bilateral infusions of muscimol into the prelimbic and infralimbiccortices in the caudal half of the MFC had no effect on sucrose pellet consumption, providing further evidence for a distinction between the role of rostral and caudal MFC in feeding. Together, these findings suggest that the adjacent medial orbital and prelimbic areas have distinct roles in consummatory behavior.

A handful of studies have investigated the role of the opioid system of the MFC in the regulation of consummatory behaviors (Selleck and Baldo, 2017). The MFC is dense with receptors for the mu opioid system (Lewis et al., 1983), and pharmacological manipulations of these cortical opioid receptors with the selective mu opioid agonist DAMGO have suggested that specific regions of the MFC and nearby OFC play a role in regulating feeding (Mena et al., 2011, 2013; Selleck et al., 2018; Giacomini et al., 202). For example, Mena et al. (2011) demonstrated that infusions of DAMGO in ventral prelimbic and lateral OFC areas potentiate feeding and lead to increases in total food intake; number of eating bouts (periods of sustained eating); and reduces the duration of these eating bouts; these measures suggest there’s an overall enhancement of the positive hedonic, or ‘liking’ of food (Castro and Berridge, 2017). Mena et al. (2013) further showed these effects are replicated with infusions of DAMGO into the infralimbic cortex and may be mediated by glutamatergic connections to subregions of the hypothalamus. Most recently, Giancomi et al. (2022) reported that reversible inactivation of the caudal prelimbic and infralimbic cortices have inverse effects on the potentiating effects of DAMGO in the agranular insular cortex. Inactivation of the infralimbic cortex positively augmented effects of DAMGO and, by contrast, inactivation of the prelimbic cortex reduced effects of DAMGO in the agranular insular cortex. The sites studied by Giancomi were more than 1 mm caudal to the focus of the present study. Together, these results support the idea that, despite falling under the umbrella of a single term “medial frontal cortex”, subregions of MFC across a rostral/caudal and dorsal/ventral axis contribute differently to reward guided consummatory behavior. and further support the heterogeneity.

Our results following infusions of DAMGO into the rostral MFC are distinct from all of these published studies. The lack of effects following DAMGO in the rostral MFC (rostral prelimbic cortex) might reflect differences in the concentration of mu opioid receptors, described as “hot spots” by Castro and Berridge (2017), in the rostral MFC and other regions such as the more caudal or lateral regions studied by Mena, Giancomi, and colleagues. It is also possible that our infusions crossed “hot spots” leading to an unclear modulation of behavior. In any case, there have been only a few studies on cortical mu receptors and their role in the control of behavior, and the findings across these studies and the present one suggest that mu opioid receptors have distinct effects in different parts of the rostral frontal cortex. Notably, a recent study on synaptic transmission by the mu opioid system reported distinct effects of DAMGO on GABA signaling in the medial and lateral OFC (Lau et al., 2020). More studies on this neurotransmitter system, which is crucial for understanding opioid abuse and addiction, are needed.

## Acknowledgements

We thank Drs Linda Amarante, David Kearns, and Jibran Khokhar for helpful comments on the manuscript.

